# Design of Transcranial Magnetic Stimulation Coils with Optimal Trade-off between Depth, Focality, and Energy

**DOI:** 10.1101/300616

**Authors:** Luis J. Gomez, Stefan M. Goetz, Angel V. Peterchev

## Abstract

**Background:** Transcranial magnetic stimulation (TMS) is a noninvasive brain stimulation technique used for research and clinical applications. Existent TMS coils are limited in their precision of spatial targeting (focality), especially for deeper targets.

**Objective:** This paper presents a methodology for designing TMS coils to achieve optimal trade-off between the depth and focality of the induced electric field (E-field), as well as the energy required by the coil.

**Methods:** A multi-objective optimization technique is used for computationally designing TMS coils that achieve optimal trade-offs between stimulation focality, depth, and energy (fdTMS coils). The fdTMS coil winding(s) maximize focality (minimize stimulated volume) while reaching a target at a specified depth and not exceeding predefined peak E-field strength and required coil energy. Spherical and MRI-derived head models are used to compute the fundamental depth–focality trade-off as well as focality–energy trade-offs for specific target depths.

**Results:** Across stimulation target depths of 1.0–3.4 cm from the brain surface, the stimulated volume can be theoretically decreased by 42%–55% compared to existing TMS coil designs. The stimulated volume of a figure-8 coil can be decreased by 36%, 44%, or 46%, for matched, doubled, or quadrupled energy. For matched focality and energy, the depth of a figure-8 coil can be increased by 22%.

**Conclusion:** Computational design of TMS coils could enable more selective targeting of the induced E-field. The presented results appear to be the first significant advancement in the depth–focality trade-off of TMS coils since the introduction of the figure-8 coil three decades ago, and likely represent the fundamental physical limit.

**DECLARATION OF INTEREST:** The fdTMS technology described in this paper is subject to a provisional patent application by Duke University with the authors as inventors. Additionally: L.J.G. is inventor on a patent pertaining to the design of focal multicoil TMS systems. S.M.G. is inventor on patents and patent applications; he has received royalties from Rogue Research, TU Muenchen, and Porsche; furthermore, he has been provided with research support and patent fee reimbursement from Magstim Co. A.V.P. is inventor on patents and patent applications, and has received research and travel support as well as patent royalties from Rogue Research; research and travel support, consulting fees, as well as equipment loan from Tal Medical / Neurex; patent application and research support from Magstim; and equipment loans from MagVenture, all related to technology for transcranial magnetic stimulation.

## INTRODUCTION

Transcranial magnetic stimulation (TMS) is a noninvasive technique using strong brief magnetic pulses that induce an electric field (E-field) in the brain, which in turn elicits or modulates neural activity. TMS is widely used in the neurosciences as a tool for probing and manipulating brain function and connectivity. Moreover, TMS is FDA-approved for the treatment of depression [1] and migraines [2] as well as for pre-surgical cortical mapping[3], and is under study for many other psychiatric and neurological disorders.

Increasing the focality and depth of TMS could enable more flexible and selective targeting of its effects. The figure-8 coil was introduced three decades ago to improve focality over single circular coils [4], and has been the standard choice for focal TMS. The literature abounds with studies of coil designs attempting further improvements in focus and penetration depth [5-19]. However, in simulation studies of a large number of existing or proposed coil designs, we and other groups showed that the designs do not exceed the depth–focality trade-off of figure-8-type coils [20]. Notably, these coil topologies were derived either empirically or by simple heuristics, reflecting the long-standing approach to TMS coil design, and suggesting that they may not be fully optimized. Therefore, an outstanding question is whether the depth–focality trade-off associated with figure-8-type coils is a fundamental physical limit, or there exist other coil designs with superior performance.

To achieve optimal depth–focality trade-off, we previously proposed a genetic-algorithm-based optimization framework applied to an array of small coils [21]. However, this computational approach did not consider all relevant energy and implementation constraints, limiting the feasibility and practicality of the resultant designs. Since TMS relies on weak inductive coupling between the coil and the brain, the coils require high-energy current pulses to induce stimulation [22]. Recent computational optimization studies demonstrated that the energy required by TMS coils can be reduced compared to conventional coils while staying within prescribed E-field characteristics [12, 23-25]. These approaches, however, do not explicitly improve the trade-off between focality and stimulation depth. Therefore, a second important question concerns the energy cost of achieving better depth–focality performance for TMS coils.

This paper proposes a methodology for designing TMS coils that achieve optimal trade-off between depth, focality, and energy of stimulation (fdTMS coils). First, a novel optimization procedure is used to determine optimum surface current distributions. Then, the surface current distributions are approximated by windings by a process that does not deteriorate their performance [12, 23, 26]. The fundamental limits of the focality vs. depth of stimulation trade-offs in a spherical head model are determined. Furthermore, energy vs. focality relations for targeting fixed depths are given. Finally, the methodology is used to determine optimal coils for targeting in a three-layer MRI derived model of the head.

## METHODS AND MATERIALS

This section details the proposed approach for designing fdTMS coils. It starts with definitions of the computational optimization problem, coil performance metrics, and parametrization of the coil current distribution. Then, methods to compute the E-field and performance metrics are described. This is followed by an optimization procedure that generates coil current distributions that achieve optimal trade-offs between stimulation depth, volume, and energy (i.e. Pareto-optimal designs). The section concludes with a method to convert the optimal current distributions to realizable coil structures, as well as a description of head models used in the simulations.

### Problem definition

Consider current distributions **I** residing on a surface *Ω* (figure 1(a)). The center **r_c_** of surface *Ω* is directly above the scalp [27] with its center surface normal **n̂_c_** oriented towards the brain (figure 1(a)). Each current distribution has identical temporal variation *p*(*t*), and a spatial variation that can be written as a linear combination of *N* modes

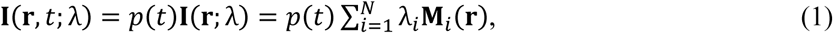

where **r** = (*x*, *y*, *z*) denotes Cartesian position, each **M**_*i*_ (**r**) is a single mode, *λ* = (*λ*_1_, *λ*_2_,…, *λ_N_*) is a vector of weights, each *λ_i_* (where *i* = 1,2,…, *N*) is a real number, and *p*(*t*) = sin(*ωt*) and *ω* = 3000 · 2*π*. Note that *p*(*t*) was assumed to be time-harmonic to simplify the exposition; however, because of the relatively low-frequency content of TMS pulses, the results apply to other current waveforms as well. The surface current modes form a vector space over the field of the real numbers with the physical character of normalized basic current distributions on a given surface on which the optimized coil windings should reside and the current **I**(**r**, *t*; *λ*) is in the span of the modes **M**_*i*_(**r**) (where *i* = 1,2,…,*N*). For example, figure 1(b) depicts three modes and a surface current resulting from a linear combination of them. During each Pareto-optimization the objective is to find weights *λ_opt_* with corresponding current distributions **I**(**r**, *t*;*λ_opt_*), termed Pareto-optimal, that achieve optimal trade-offs with respect to a combination of performance metrics. In this manuscript we consider the following three metrics:

**Figure 1.**
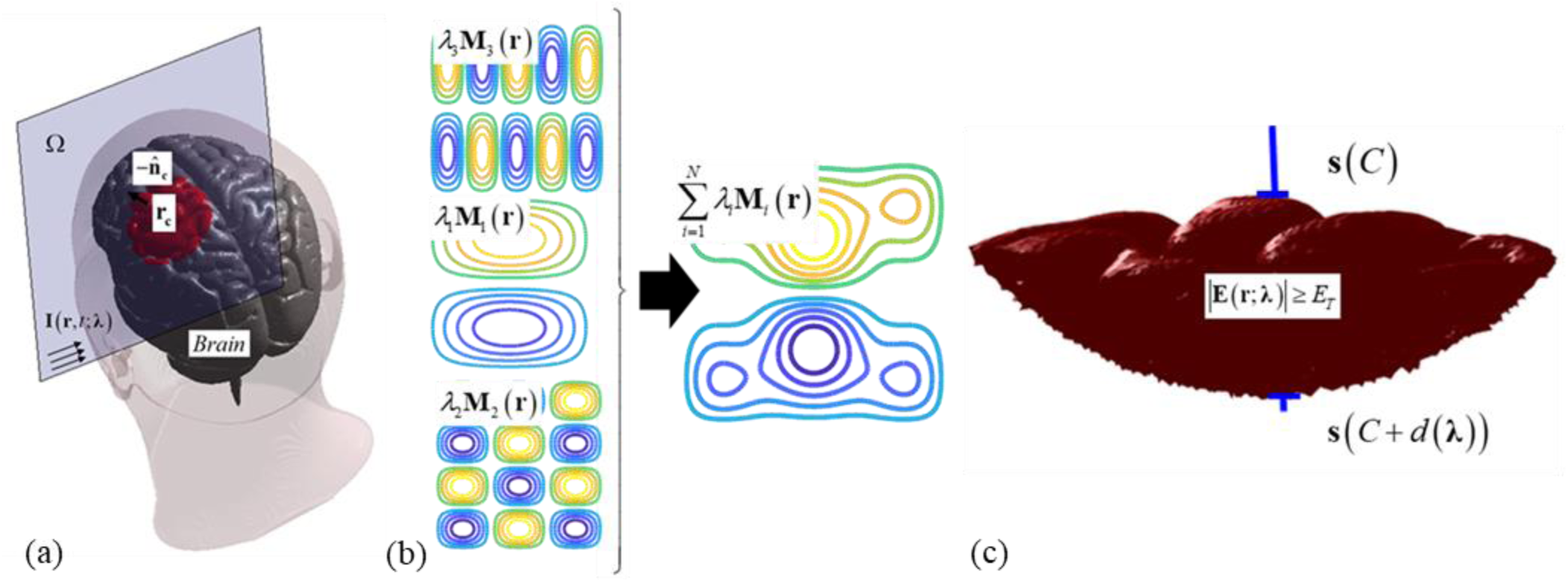
(a) Example coil current having a square support surface and placed directly above the scalp. (b) Contours of three current modes and the coil surface current resulting from a linear combination of them. (c) The stimulated region depicted in red pierced by a blue column along which depth is measured.

i. *Minimum stimulation volume*: Consistent with previous TMS coil design studies, it is assumed that brain tissue that is exposed to a peak E-field above *E*_TH_ = 50 V/m is stimulated [20]. Note that due to the linearity of the problem the specific choice of this E-field threshold does not influence the depth, focality, and energy optimization; it only affects the absolute energy. The stimulated volume *V* is defined as

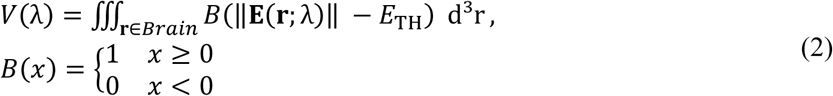

where **E**(**r**; *λ*) denotes peak E-field at location **r** induced by the surface current **I**(**r**, *t*; *λ*), ∥·∥ denotes vector magnitude, *B*(*x*) is a unit step function, and the integration is over the brain region (denoted *Brain*). *B*(∥**E**(**r**; *λ*)∥ − *E_TH_*) indicates whether ∥**E**(**r**;*λ*)∥ is above the stimulation threshold. For example, in figure 1(a),(c) the stimulated region is depicted in red and its volume is equal to *V* (*λ*).
ii. *Maximum depth of stimulation:* At the targeted depth *d* it is desired to preferentially stimulate neuronal elements aligned along a direction **t̂**. As such, the E-field magnitude along **t̂** must equal or exceed the stimulation threshold. The depth of stimulation *d* is defined along a line ***s***(*l*) chosen as a line that intersects at and is perpendicular to the center of the surface current support (i.e. ***s***(*l*) = **r_c_** + *l***n̂_c_**). In accordance, the stimulation depth is

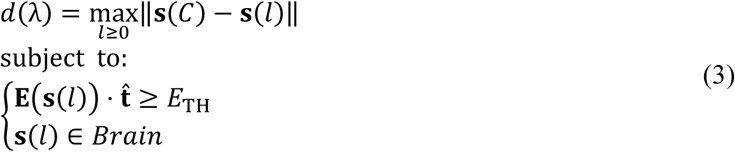

where ***s***(*C*) denotes the point on the cortex closest to **r_c_**. Figure 1(c) depicts the stimulated region from figure 1(a). The line ***s***(*l*) that traverses the brain is shown in blue along with markers at positions ***s***(*C*) and ***s***(*C* + *d*(*λ*)); *d*(*λ*) is the distance between these two markers. (Note that the choice of **n̂_c_** pointing toward the brain results in ***s***(*C* + *d*(*λ*)) being the deepest point stimulated.)
iii. *Minimum energy:* TMS pulses have relatively low-frequency temporal variation and their induced magnetic field is negligibly affected by the presence of the head. The magnetic energy stored in the current distribution can be computed using the Biot-Savart law [28] as

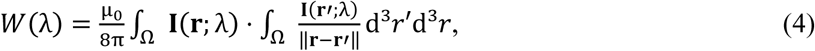

where μ_0_ is the permeability of free space.

Apart from the above metrics, we combine the *V* and *d* to define spread as the average transverse surface area of the stimulated region, *S* = *V/d* [20, 29]. A decrease in *V* or *S* is equivalent to an increase in focality. Furthermore, safety considerations limit the peak E-field that brain tissue can be exposed to. For a given *α*, we assume that E-field strengths exceeding *αE_TH_* in the brain are unacceptable. Therefore, currents in the span of the modes that result in an E-field that exceeds *αE_TH_* in the brain are excluded from the admissible designs. In this work, we arbitrarily set *α* = 2 which results in *V* defined as the subvolume of the brain where the E-field equals or exceeds half of its peak value, *V*_1/2_, *d* defined as the largest depth where the E-field equals or exceeds ½ of its peak value, *d* = *d*_1/2_, and *S* = *S*_1/2_ = *V*_1/2_/*d*_1/2_; these metrics are commonly used to characterize TMS coils [20, 29, 30].

### Computation of performance metrics

Execution of the Pareto-optimization procedure to determine optimal trade-offs between metrics (i)-(iii) requires evaluations of the E-field **E**(**r**, *t*; *λ*), stimulated depth *d*(*λ*), and stimulated volume *V*(*λ*) induced by currents **I**(**r**; *λ*), as well as their energy *W*(*λ*) for hundreds of thousands of values of *λ*. This section, therefore, describes a technique for computing these quantities rapidly.

The head tissues have approximately constant conductivities at TMS frequencies and are non-magnetic. Consequently, the E-field **E**(**r**, *t*;*λ*) is a linear combination of fields due to *p*(*t*)**M**_*i*_(**r**) (where *i* = 1,2,…, *N*). Furthermore, because TMS pulses are relatively low-frequency, the spatial and temporal variations of the E-field are separable. Thus,

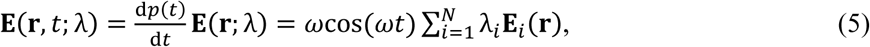

where **E**_*i*_(**r**) is the peak E-field due to a single mode current *p*(*t*)**M**_*i*_(**r**). E-fields **E**_*i*_(**r**) are determined using a method described in [31]. The total E-field induced in the head is the sum of the primary E-field due to the surface current and a secondary contribution from the scalar potential, –∇ϕ,

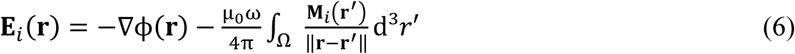

and

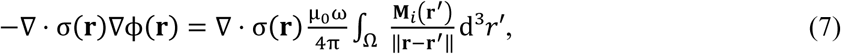

where σ(**r**) is the tissue conductivity, and *n̂* · **E**_*i*_(**r**) = 0 on the surface of the scalp. To solve for ϕ and ∇ϕ, first, the head model is discretized into a tetrahedral mesh having *P* nodes, *Q* edges and each tetrahedron is assigned a constant tissue conductivity. The scalar potential ϕ is approximated by piece-wise quadratic nodal elements *N_m_*(**r**) (where *m* = 1,2,…,*P* + *Q*) [32]. Weak forms of Eq. (7) are sampled also using piecewise-quadratic nodal elements as testing functions in a standard Galerkin procedure. This results in

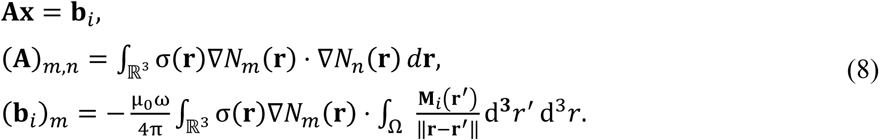

Entries (**A**)_*m*,*n*_ are computed analytically using expressions provided in [32]. To determine (**b**_*i*_)_*m*_, the outer integral is approximated using a midpoint rule and the inner-integration is done by discretizing the surface current support *Ω* into a surface triangle mesh and assuming the current is constant within each triangle. Then, the inner-integration over each triangle is done analytically using expressions in [33]. The system of equations (8) is solved using a transpose-free quasi-minimal residual [34] iterative solver to a relative residual of 10^−6^.

Computing *d*(*λ*) and whether the current distribution induces field strengths that exceed the safety threshold can be trivially done by using the E-field. Furthermore, the E-field sampled on mesh tetrahedron centroids in the brain is used to determine the value of *V*(*λ*) numerically. The explicit formula used for approximating the stimulated volume *V*(*λ*) is

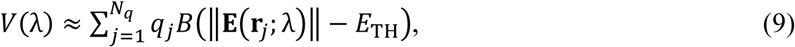

where *N_q_* is the number of sample points for the approximation, and *q_j_* and **r**_*j*_ are the volume and centroid location of the *j*-th tetrahedron, respectively.

Equation (10)

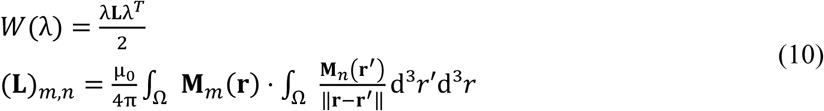

is used to determine the energy. Entries (**L**)_*m*,*n*_ are determined by discretizing the surface current support Ω into triangles and assuming that the mode function is constant within each triangle. A 200^th^ order accurate Gauss quadrature rule is used for numerical integration of the outer integral, and the inner integral is computed exactly using formulas in [33]. (Note that the E-fields **E**_*i*_(**r**) and mutual inductance matrix **L** are precomputed prior to executing the optimization; this enables rapid evaluation of *V*(*λ*), *d*(*λ*), and *W*(*λ*).)

*Mixed integer linear programming algorithm for finding optimal surface currents*

Next, we define *λ′* _*opt*_(*Ṽ*, *d̃*) as

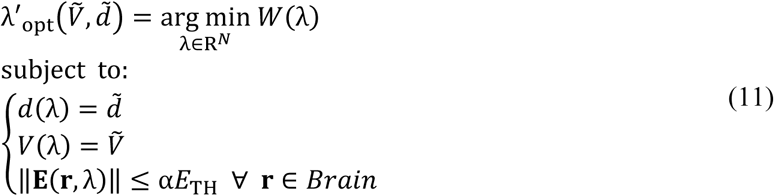

and

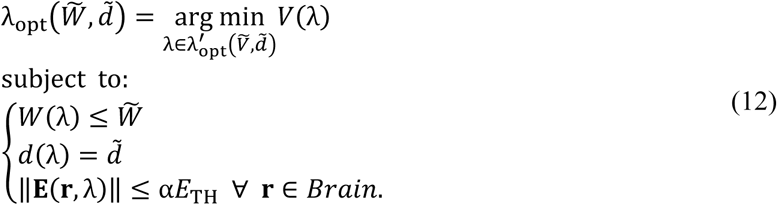

The current **I** (**r**; *λ′* _*opt*_(*Ṽ*, *d̃*)) minimizes the energy to achieve a given combination of *V*(*λ*) = *Ṽ* and *d*(*λ*) = *d̃* while not exceeding the E-field safety threshold. Note that many combinations of *Ṽ* and *d̃* are not physically achievable and they are not in the domain of *λ′* _opt_(*Ṽ*, *d̃*). The current distribution **I**(**r**; *λ*_opt_(*W͠*, *d̃*)) = **I**_opt_(**r**; *W͠*, *d̃*) minimizes (and requires at most *W͠*) energy to induce fields that stimulate up to a *d̃* depth into the brain while minimizing the stimulated volume. The Pareto-optimal current distribution **I**_opt_(**r**; *W͠*, *d̃*) achieves optimal trade-offs between stimulation depth, volume, and energy.

The Pareto function *V* (*λ*_opt_(*W͠*, *d̃*)) = *V*_min_(*W͠*, *d̃*), provides the smallest stimulated volume possible while reaching a given target depth *d̃* and not exceeding energy *W͠*. Quantity *V*_min_(*W͠*, *d̃*) and additional Pareto functions

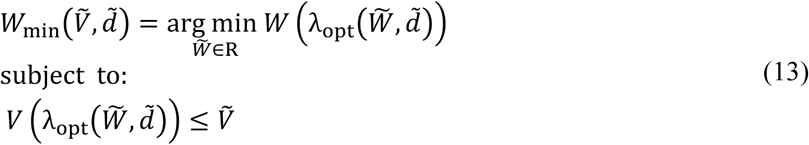

and

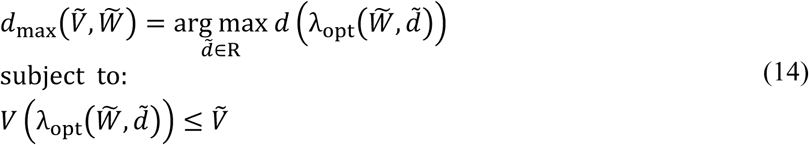

are the optimal achievable trade-offs between the three objectives. To determine *λ*_opt_(*W͠*, *d̃*), *V*_min_(*W͠*, *d̃*), *W*_min_(*Ṽ*, *d̃*), and *d*_max_(*Ṽ*, *W͠*) starting from *d̃* = *d*_start_ and *W͠* = *W*_start_, and slowly incrementing *d̃* and *W͠* to *d̃* = *d*_end_ and *W͠* = *W*_end_, respectively, we carry out a sequence of optimizations, each solving Eq. (12) (detailed below).

Computation of the cost *V*(*λ*) using Eq. (9) involves a sum of unit step functions. Step functions are convex for argument less than zero and concave for arguments more than zero. As a result, the number of inflection points increases with increasing number of summands (i.e. increasing *N*_q_) and the optimization problem in Eq. (12) is highly non-convex. In the optimization literature problems of the form of Eq. (12) are known as sigmoidal programming and their complexity is NP-hard with respect to the number of inflection points. For many practical problems, Eq. (12) can be solved using branch and bound algorithms that branch at the inflection points of the cost [35]. Guided by insights from [35], we discretize the constraints and add slack variables to convert Eq. (12) into a Mixed-integer linear program (MILP) that can be solved using Matlab’s branch and bound MILP solver [36]. Furthermore, to minimize *N_q_* the optimization is done iteratively by performing a series of MILP optimizations each time adaptively refining the sample point locations by a scheme detailed in the Adaptive Refinement section of the Appendix.

The non-convex objective is converted to a linear one by using binary slack variables *t_j_* ∈ {0,1} (where *j* = 1, 2,…, *N*_q_) that encode the inflection points of the cost function. The inequality constraints of Eq. (12), which enforce the initially defined safety limit for the coil, are replaced with the following

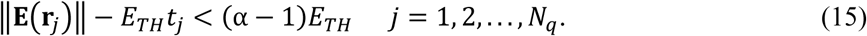

Constraints of the form of Eq. (15) ensure that if *t_j_* = 0, location **r**_*j*_ is not activated (i.e. |**E**(**r**_*j*_)| < *E_TH_*) and if *t_j_* = 1 the E-field does not exceed the safety threshold (i.e. ∥**E**(**r**_*j*_)∥ < *αE*_TH_). Correspondingly, the optimization problem in Eq. (12) becomes

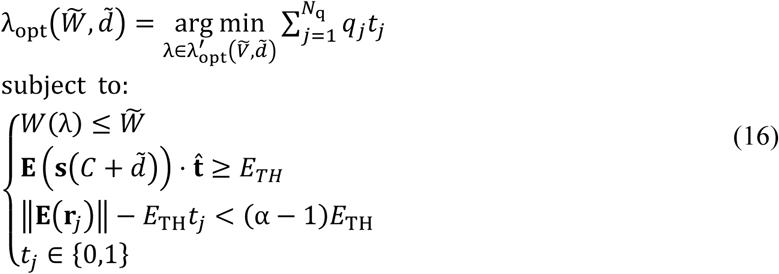

Optimization problem in Eq. (16) now has a simple linear cost function. All of the complexity of Eq. (12) is now encapsulated in the binary slack variables, which separate the convex regions (*t_j_* is zero) and concave regions (*t_j_* is one) of the summands of Eq. (9). More importantly, this choice of slack variables will result in branching at inflection points for the branch and bound algorithm, thereby, enabling its fast convergence.

We have a number of nonlinear constraints that limit the E-field magnitude and energy. The nonlinear constraints can be approximated arbitrarily accurately by linear ones. Here this is done to lower computational costs of the optimization. Like in [23], each of the nonlinear E-field magnitude constraints of Eq. (15) are approximated by 162 linear constraints:

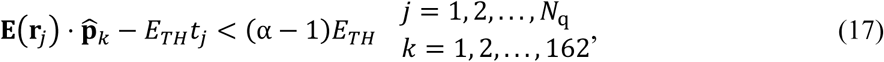

where **p̂**_*k*_ is a unit vector along a single direction. Each of the constraints in Eq. (17) are of the same form as Eq. (15), with the magnitude of the electric field replaced with the magnitude along a single predetermined direction. The 162 predetermined directions are chosen to uniformly and densely span a unit sphere to approximate all possible directions of the E-field. The predetermined directions are chosen as the locations of vertices of a twice-barycentrically-refined regular icosahedron that is centered about the origin and projected onto a unit sphere as described in [37]. The resultant directions guarantee that one of the projections has a magnitude that is within 1.78% of the true E-field magnitude. Correspondingly, satisfying all constraints of Eq. (17) guarantees that the constraints of Eq. (15) are satisfied with a maximum error of 1.0%. For illustration, a 2D example of this procedure is shown in figure 2(a). The E-field vector is depicted in red and directions vectors **p̂**_1_ to **p̂**_8_ are chosen each pointing toward a single vertex of the green octagon. The best estimate of |**E**(**r**_*j*_)| is **E**(**r**_*j*_) · **p̂**_3_ and its error corresponds to the distance along the direction of **E**(**r**_*j*_) between the green octahedron and the red circle.

**Figure 2.**
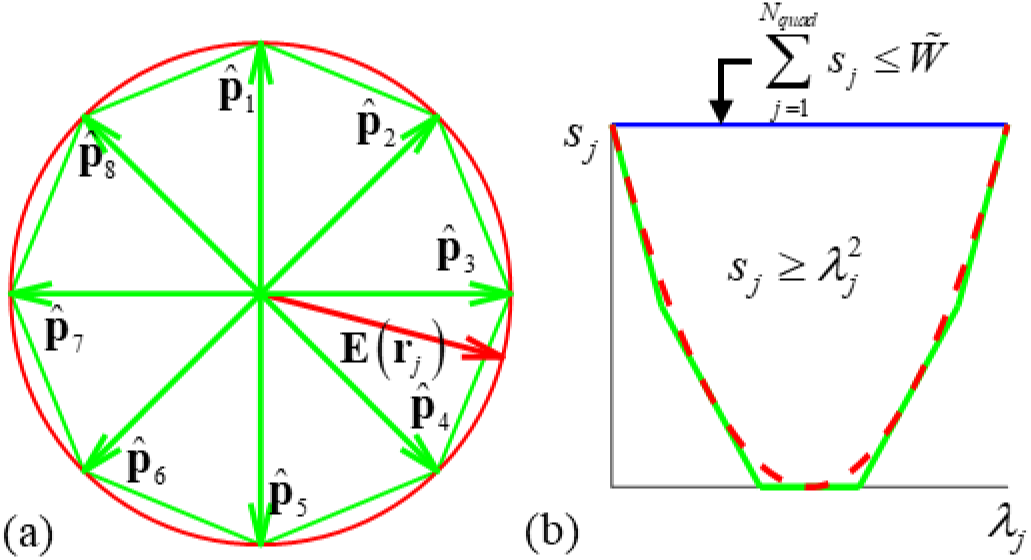
(a) Example of the E-field magnitude estimation process: Unit E-field (red) on the plane is projected onto all of unit vectors **p̂**_1_ to **p̂**_8_. Since constraints are applied to all projections, the maximum of all projections is the E-field magnitude estimate. (b) Example of the approximation of energy constraints. The feasible region for *s_j_* is above red the dashed curve and below the blue line; its piece-wise linear approximation is shown with green lines.

The quadratic constraint *W*(*λ*) ≤ *W͠* can also be approximated by a number of linear ones. To do this, we first assume that the modes are energy orthonormal. (If the modes are not orthonormal, they need to be converted to orthonormal ones; the approach to do so used here is given in the Mode Preprocessing section of the Appendix.) Energy orthonormal modes have the property

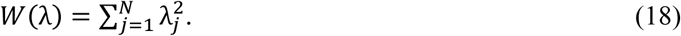

In other words, their mutual inductance matrix **L** is diagonal with twos along its diagonal. First, we introduce slack variables and slack variable constraints:

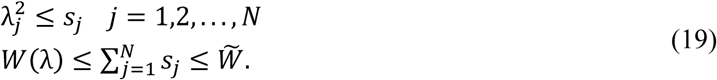

The linear inequality constraint on the second line of Eq. (19) ensures that energy is bounded from above by *W͠*. We replace each quadratic constraint of the form
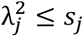
with *N*_Δ_ linear ones. A linear estimate of *λ_j_* ^2^ that coincides with *λ_j_* ^2^ at *x* (i.e. *λ_j_* ^2^ ≈ − *x*^2^ + 2*xλ_j_*) will always underestimate *λ_j_* ^2^ by an amount that grows quadratically away from *x* as (*λ_j_* − *x*)^2^. On the range of admissible values for
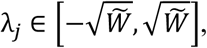
we approximate *λ_j_*^2^ as the maximum of *N*_Δ_ linear approximations of *λ_j_*^2^ each coinciding with *λ_j_*^2^ at points spaced Δ apart; this will result in a maximum possible error of Δ^2^/4. The resultant constraints are

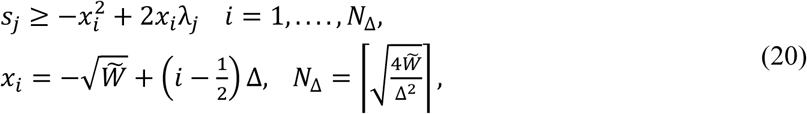

where ⌈·⌉ rounds to the nearest integer. If all constraints are satisfied, the energy will be guaranteed to be
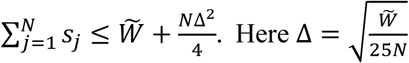
to achieve a maximum possible error of 1.0% in the energy estimate. For example, in figure 2(b) the feasible range of values for *S_j_* is depicted as the region above the red dashed curve and below the blue line. Linear constraints of form Eq. (20) with a choice of
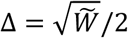
force *S_j_* to be above the green lines. In other words, the linear constraints form a piece-wise linear discretization of the quadratic constraints, and by choosing a smaller Δ we can approximate arbitrarily well the nonlinear constraint (i.e. red dashed line).

The final MILP optimization problem is

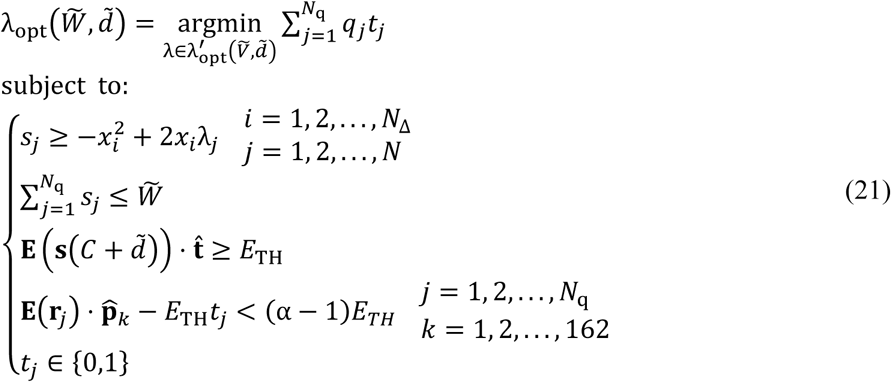

Note that in Eq. (21) energy is not optimized; it is only restricted. Improving focality requires increasing sharpness of induced cortical E-field, which results in increased energy requirements [12, 13]. As such, minimizing stimulated volume while restricting energy will result in an energy optimal design.

### Generating coil windings

The above procedure yields Pareto-optimal surface currents **I**_*opt*_(**r**; *W͠*, *d̃*) for penetration depths *d*_*start*_ < *d* ≤ *d*_end_ and energy levels *W*_start_ ≤ *W͠* ≤ *W_end_*. However, practical implementation of fdTMS coils requires the use of coil windings that can be driven by a TMS coil driver. Here the continuous surface current distributions are transformed into separate coil windings by a procedure originally developed for deriving MRI gradient coils from ideal continuous current distributions [26] and more recently used to design minimum energy TMS coils [12, 25]. In summary, surface currents are replaced by a design having at most *N_t_*/2 concentric turns. This is done by tracing out contours of its stream function *Sr*(**I**(**r**; *λ*)). The contours levels *L_k_* (where *k* = 1,2,…,*N_t_*) are chosen as

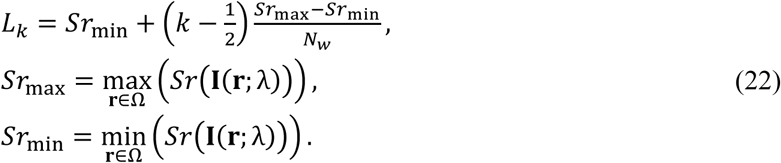

The turns resulting from tracing out the contours are connected serially by a feed that minimally affects radiation and thereby performance. This procedure produces windings that, for a large enough *N_t_*, can match the magnetic moment of the continuous current distribution to a prescribed accuracy. More importantly, even for relatively low-values of *N_t_*, the resultant coils generate fields that match those of the original surface current distributions in the brain. Here, *N_t_* is chosen to be large enough to both achieve the same E-field in the brain and a target inductance between 9 and 15 μH to be compatible with existing TMS driving sources.

### Coil support surfaces

The procedure described above is used to design fdTMS coils with either a sphere shell, hemi-sphere shell, or square planar support.

#### Sphere shell

The sphere shell support is centered about the origin, has a radius of *r*_0_ = 9 cm, and its modes **M**_*l*,*m*_ are chosen as [12]

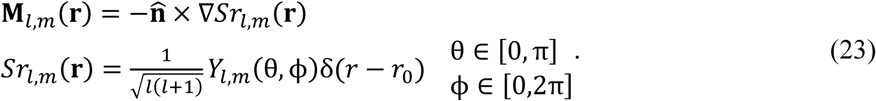

Here **r** = (*r*, θ, ϕ) is in spherical coordinates, the unit normal points in the radial direction (i.e. **n̂** = **r̂**), *Sr*_*l*,*m*_(**r**) is the stream function corresponding to mode (*l*, *m*), *Y*_*l*,*m*_ is the spherical harmonic with normalization constants [12], and δ(·) is a delta function. To compute the spherical harmonics accurately we use stable recursion relations [38]. The spherical surface current is centered about the spherical head, as depicted in figure 3(a).

**Figure 3.**
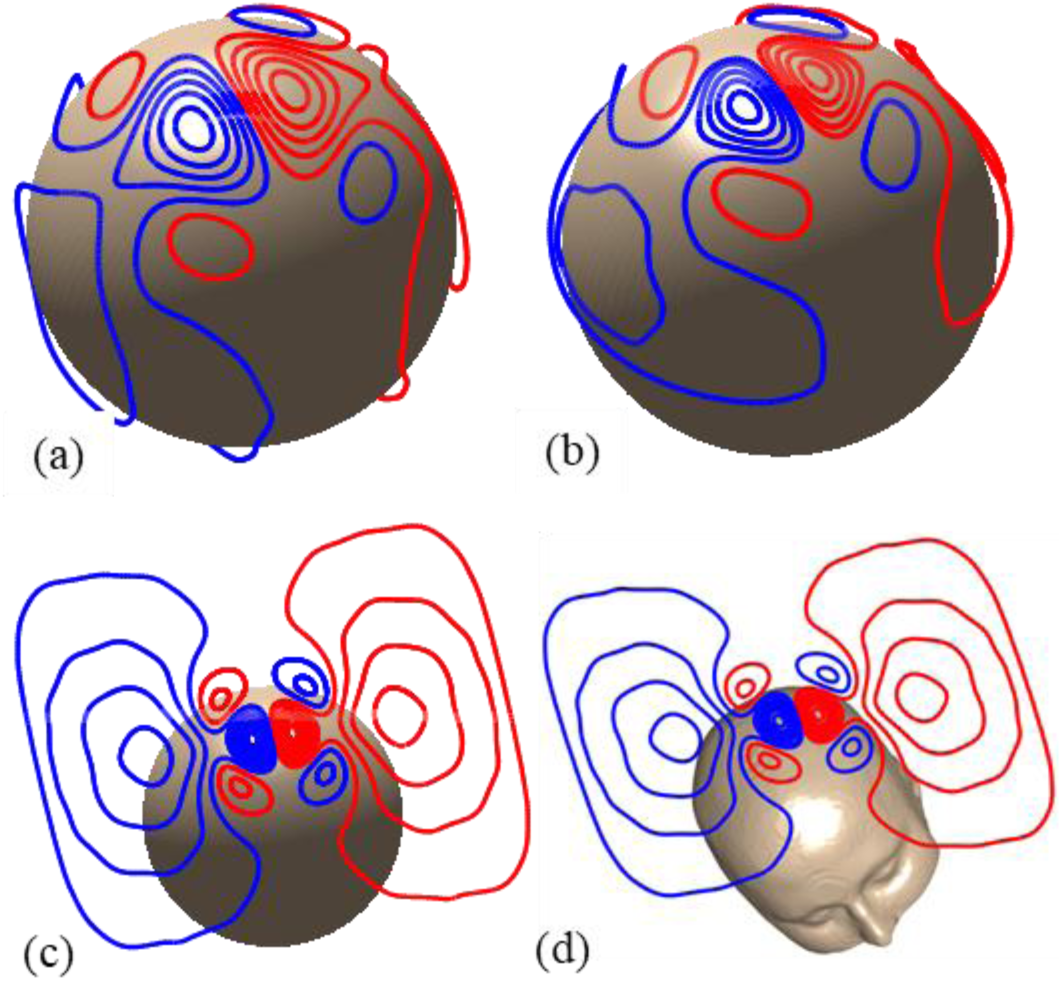
Example fdTMS coil designs. (a) Spherical coil surface with radius 9 cm and concentric with the spherical head model. (b) Half-sphere surface coil with radius 9 cm. (c) 32 × 32 cm square coil surface centered 5 mm above the apex of the spherical head model. (d) 32 × 32 cm square coil centered 5 mm above the scalp over the brain target.

#### Half-sphere shell

The half-sphere shell also has a radius of *r*_0_ = 9 cm and its modes **M**_*l*,*m*_ are chosen as

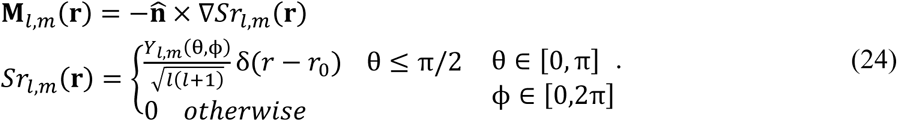

To ensure all modes form complete loops only the modes that have a zero valued stream function at the equator (i.e. *Y*_*l*,*m*_(π/2,ϕ) = 0 ∀ ϕ) are included in the optimization. The half-sphere surface current is centered about the spherical head, as depicted in figure 3(c).

#### Planar square

The surface current resides on a 32 cm × 32 cm square plane. Expressions for the modes **M**_*m*,*n*_ (**r**) and corresponding stream functions *Sr*_*m*,*n*_ (**r**) are given for a local coordinate system where the square support is assumed to reside in the region *x*,*y* ∈ [−0.16 cm,0.16 cm]

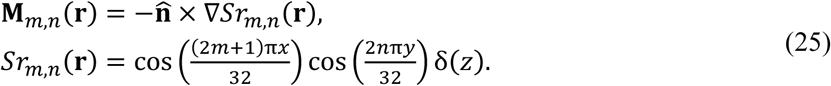

Here **n̂** = **ẑ**. To place the surface current appropriately relative to the head, the above expressions are translated and rotated with appropriate local-to-global coordinate transformations. For the sphere head simulations, the planar square support coil is placed 5 mm above the apex of the spherical head model, as depicted in figure 3(c). For MRI-derived head simulations, the coil is placed 5 mm above the scalp and directly above the hand knob region of the primary motor cortex, as depicted in figure 3(d).

### Head models

#### Spherical head model

A common sphere head model consisting of a homogenous sphere with conductivity 0.33 S/m and total radius of 8.5 cm is used. The head model consists of two concentric spheres each centered about the origin and having radii of 7.0 cm and 8.5 cm, respectively. The inner sphere corresponds to the brain, and the outer shell—to the CSF, skull, and skin. This spherical model was used to characterize various TMS coils in [20] and in optimization studies [12]. For each simulation (see figure 3(a)-(c)), the center of the coil in Cartesian coordinates is **r_c_** = (0,0,0.09) and **n̂_c_** = −**ẑ**. Correspondingly, depth is measured along − *z* direction and starting from *z* = 7.0 cm. Note that analytical expressions for the E-field generated inside the spherical head model are given in [12] and used in lieu of the FEM solver to determine the E-field generated by the surface currents.

#### MRI-derived head model

The MRI-derived head model uses the SimNIBS segmented head mesh [39]. The original tissue model consists of five tissue types: gray matter (GM), white matter (WM), CSF, skull, and scalp. We combine GM, WM, CSF into a single compartment resulting in a three layer head model [12]. The conductivities are chosen as 0.01 S/m for the skull, 0.465 S/m for the scalp, and 0.276 S/m for the intracranial space [27]. The coil coordinates on the SimNIBS mesh coordinate system are **r_c_** = (−4.0,−0.4,4.8) cm,
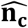
= (0.55,−0.005,−0.83), and **t̂** = (−0.027,0.99,−0.024), as depicted in figure 3(d).

## RESULTS

### Depth vs. focality relations

First, the optimization approach is used to determine the fundamental limits of focality as a function of depth of stimulation. Results are shown in figure 4 along with results from [29]. Existing coils exhibit suboptimal trade-offs between focality and depth. For fdTMS designs with *d*_1/2_ of 1.0–3.4 cm, the spread (or, equivalently, volume) can be theoretically decreased by 42%–55% compared to existing TMS coils without decrease in penetration depth. Comparisons with the Magstim 25 mm figure-8 (P/N 1165), Magstim 70 mm figure-8 (P/N 3190), and MagVenture double-cone (DB-80) commercial coils are given in Table 1. Halving of *S*_1/2_ while maintaining the stimulation depth can be achieved theoretically. Decreasing *S*_1/2_ requires inducing sharper E-fields in the cortex, which results in increased energy requirements [13, 23]. The energy required by these unconstrained-energy fdTMS coils is one to three orders of magnitude higher than conventional TMS coils, which is impractical. In the next section, we address the relationship between coil energy and focality improvement, showing that a significant proportion of these gains can be achieved with feasible energy requirements.

**Figure 4.**
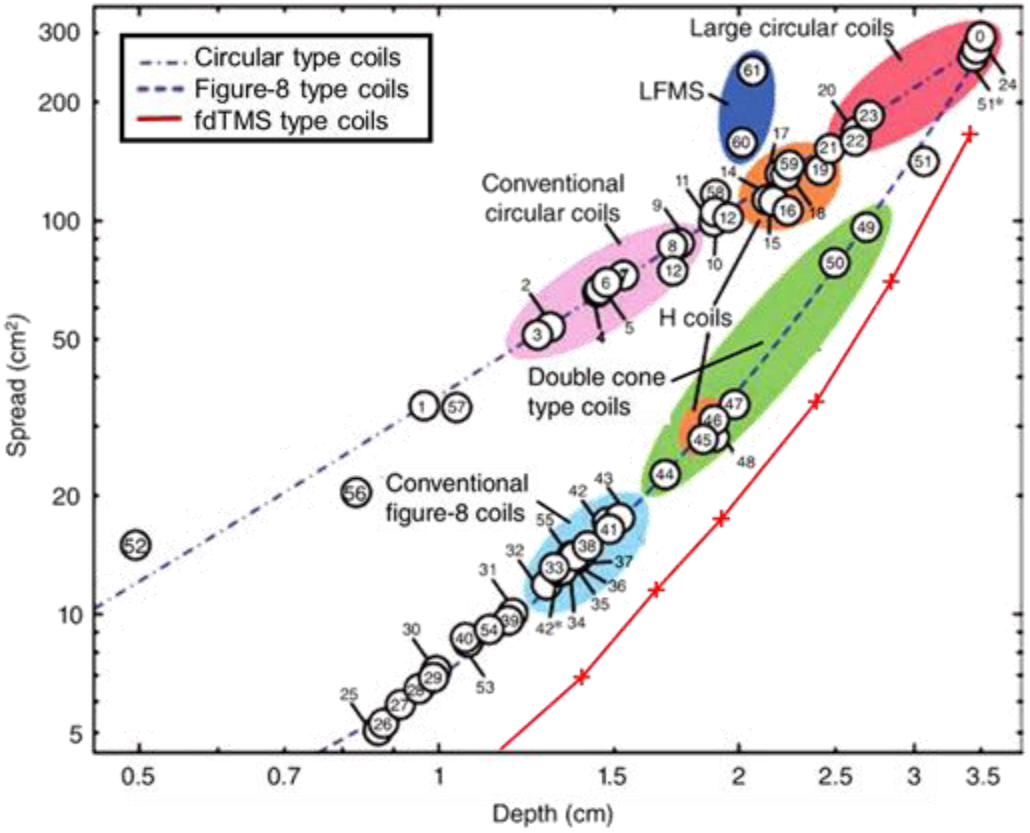
Optimal trade-off between spread, *S*_1/2_, and depth, *d*_1/2_ achieved in this study (red curve with “+” markers denoting individual designs). For comparison, the performance of other coil designs is reproduced, with permission, from [29]; see the latter reference for identification of the specific coils corresponding to the numbers.

**Table 1.** Depth *d*_1/2_ spread *S*_1/2_, and energy *W* of three conventional coils and the corresponding fundamental (unconstrained energy) limit of focality improvement achievable with fdTMS coils.

| Conventional coil |  |  |  | fdTMS focality limit |  |  |
| --- | --- | --- | --- | --- | --- | --- |
| Type | $d_{1/2}$(cm) | $S_{1/2}$ (cm<sup>2</sup>) | $W$ (J) | $d_{1/2}$ (cm) | $S_{1/2}$ (cm<sup>2</sup>) | $W$ (J) |
| 25 mm figure-8 | 1.0 | 6.5 | 212 | 1.0 | 2.8 | $74 \times 10^3$ |
| 70 mm figure-8 | 1.4 | 14.8 | 106 | 1.4 | 7.1 | 508 |
| Double cone | 1.7 | 22.6 | 57 | 1.7 | 12.0 | $11 \times 10^3$ |

### Energy vs. focality at a fixed depth

Here we analyze trade-offs between focality and energy usage for various target depths and coil topologies. The spherical head is used again. The three different coil surface types (i.e. sphere, half-sphere, and square) are placed as shown in figure 3(a)–(c). In figure 5, we show energy vs. spread curves for target depths *d*_1/2_ = {1.0, 1.4, 1.7} cm. For a fixed energy level, sphere coils are more focal than the others and half-sphere ones are more focal than square ones. This difference is more pronounced for deeper targets than for shallower ones. For example, for fdTMS designs with target depth *d*_1/2_ = 1.0 cm (figure 5(a)), the sphere and hemisphere coil performance curves appear to merge with increasing energy. For fdTMS designs with target depth *d*_1/2_ = 1.7 cm, the sphere, half-sphere, and square coils have minimum spread *S*_1/2_ equal to 12.7, 14 and 17.8 cm^2^, respectively. All fdTMS coil designs exhibit either improved focality and/or improved energy over existing designs. For the square topology, improvements over the MagVenture double cone coil are marginal, demonstrating that to target deeper into the head it is preferable to have a topology that conforms to the head (or a square coil with increased size, as discussed below).

**Figure 5.**
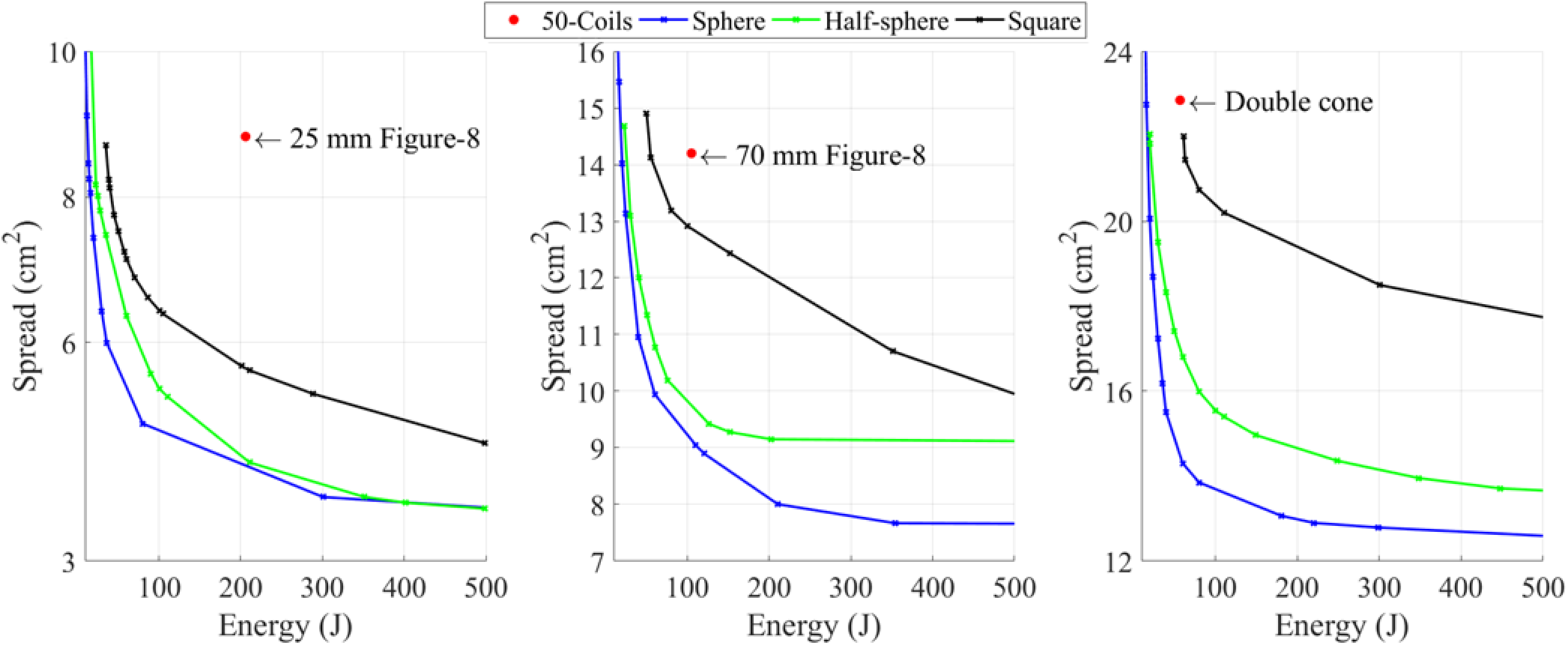
Energy, *W*, vs. spread, *S*_1/2_, curves for optimized fdTMS coils with sphere, half-sphere, or square surface for cortical-surface-to-target depths of (a) *d*_1/2_ = 10 mm, (b) *d*_1/2_ = 14 mm, and (c) *d*_1/2_ = 17 mm matching three conventional TMS coils.

In figure 6, we show fdTMS designs for targeting *d*_1/2_ = 1.4 cm into the brain, each exhibiting a different focality *S*_1/2_. Compared to a conventional figure-8 coil, the stimulated volume can be decreased by 36%, 44%, or 46%, for matched, doubled, or quadrupled energy. Unsurprisingly, the minimum energy sphere and square coils resemble the minimum energy coils of [12, 23]. With decreasing *S*_1/2_, the windings laterally concentrate and new reverse polarity windings appear (figure 6(a)–(e)). These windings partially cancel superficial fields and enable field shaping. Square coils have windings that cluster on the boundary of the square region. This suggests that making the coil of increased size could result in performance improvements. However, given the already large size of the square coil and superior performance of sphere and hemi-sphere coils, curving the square shape might be more efficient.

**Figure 6.**
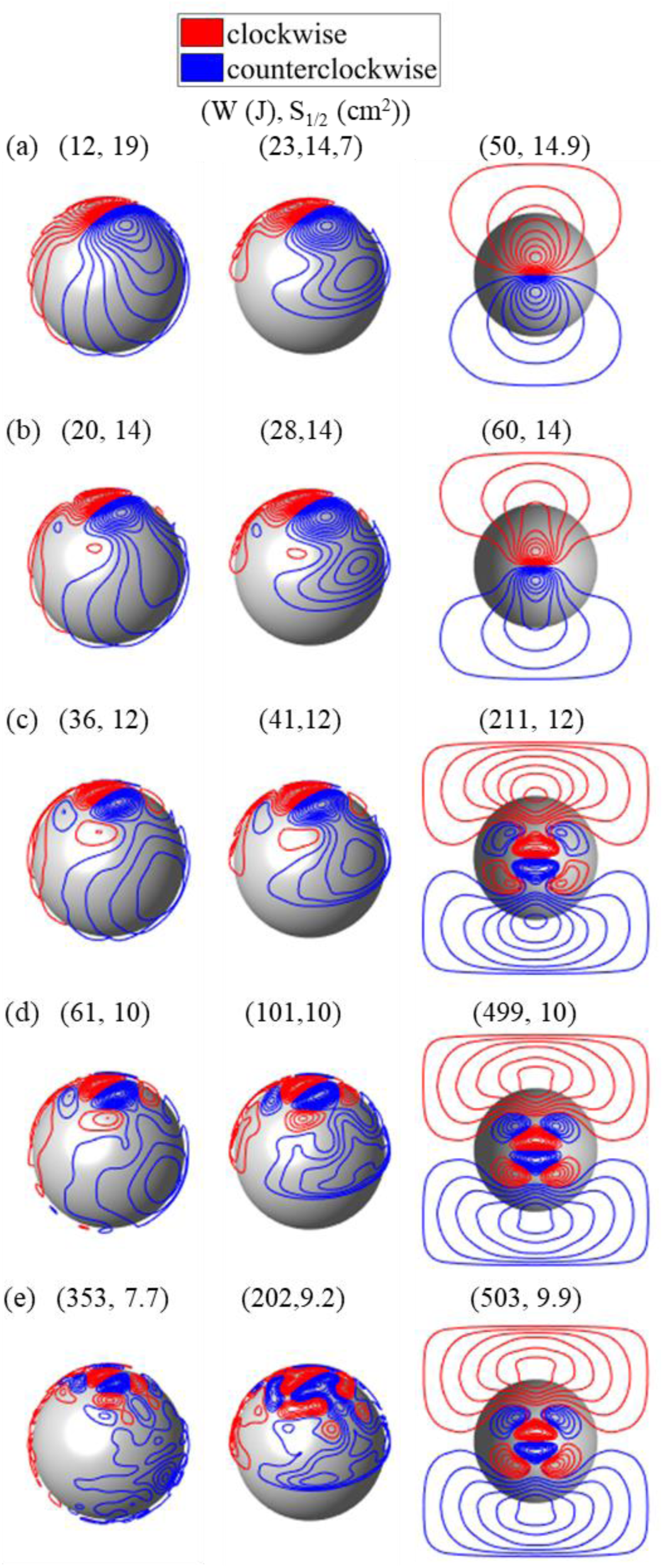
fdTMS coil designs for target depth of 14 mm, corresponding to conventional figure-8 coil. Designs with (a) minimum energy, (b) *S*_1/2_ = 14 cm^2^, (c) *S*_1/2_ = 12 cm^2^, (d) *S*_1/2_ = 10 cm^2^, and (e) minimum *S*_1/2_ (no energy constraint).

### MRI derived head model results

Here we use the optimization framework to develop designs that improve targeting in the MRI-derived head model. First, the E-field generated by a standard Magstim 70-mm-loop-diameter figure-8 coil (coil #31 in [20]) is determined. Just like the planar surface current support, the figure-8 coil is placed 5 mm above the scalp and centered directly above the hand knob region of the primary motor cortex as depicted in figure 7. We determined *S*_1/2_, *d*_1/2_, and *W* of the figure-8 coil to be 11.9 cm^2^, 1.1 cm, and 34 J, respectively. Then, we used the optimization framework to design Pareto optimal coils that have the same *d* while minimizing *V*_1/2_ (equivalently *S*_1/2_) and *W*. The resulting Pareto front is shown in figure 8. For the same energy requirements, the spread can be decreased by 16% compared to the figure-8 coil. Like the energy optimal coils of [23] for the same spread we observed that the energy can be lowered by 38%. By doubling and quadrupling the energy, the spread can be decreased by 27% and 37%, respectively.

**Figure 7.**
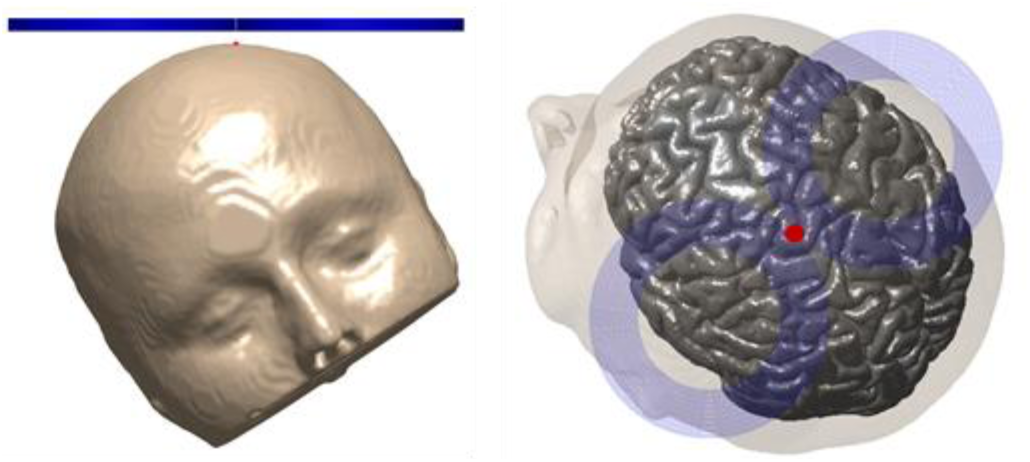
Figure-8 coil setup (a) front view (b) top view. The coil is centered (red dot) above the primary motor cortex hand knob.

Figure 9 shows the coil windings, E-fields distributions, and stimulated region on the brain surface for various coil designs along with the Magstim figure-8 coil results. We observe similar winding patterns as those of the previous section; however, the actual windings have different sizes. Here the individual windings are more circular and smaller relative to those designed for the spherical head. Upon inspection of the E-field and stimulated region maps, it is evident that with increasing energy the fields become more focal.

**Figure 8.**
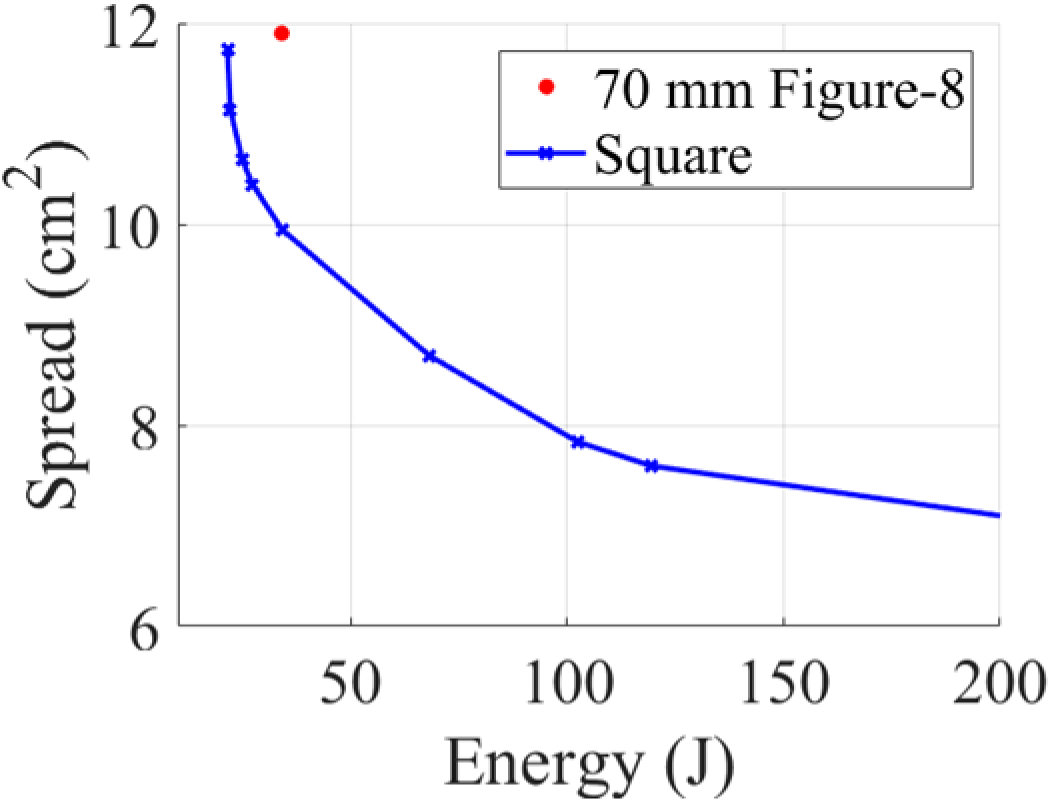
Energy, *W*, vs. spread, *S*_1/2_, curves for hand knob stimulation for a conventional figure-8 coil and optimized fdTMS coil on a 32 × 32 cm square plane.

**Figure 9.**
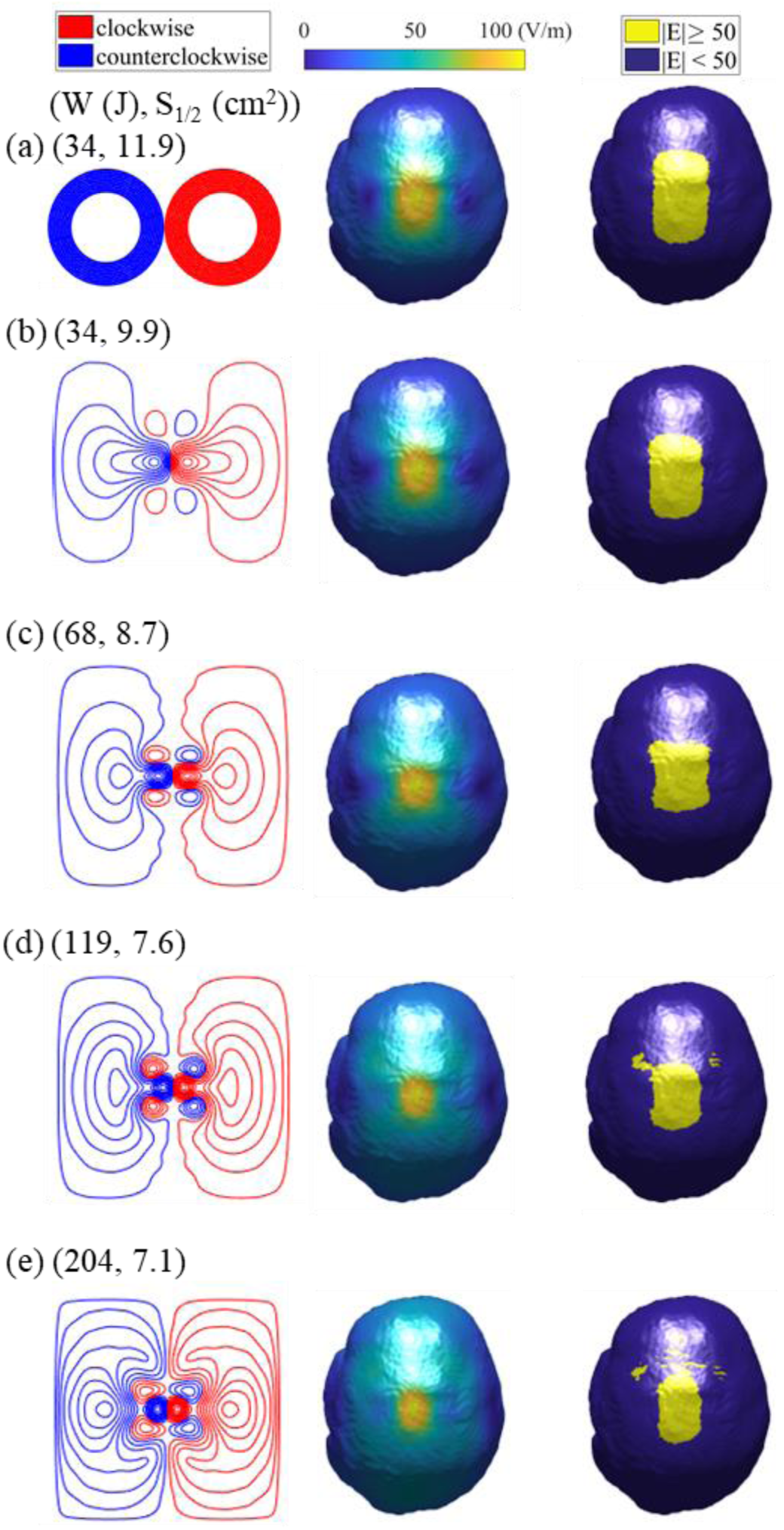
Coil designs (left), E-field distributions in the brain (middle), and stimulated brain regions (right) for (a) Magstim figure-8 and optimized fdTMS coils with energy (b) *W* = 34 J, (c) *W* = 68 J, (d) *W* = 120 J, and (e) *W* = 200 J.

## DISCUSSION

We developed and applied a method for designing TMS coils that achieves optimal trade-offs between depth, focality, and energy. By enabling control of the energy requirements of the coil, the methodology can be used to design fdTMS coils that are feasible and compatible with any coil driver. Furthermore, the methodology is general and can be applied to various coil surfaces and head models. On a triangle mesh approximation of an arbitrary coil surface, modes can be assigned as loop-finite elements on vertices of the mesh [40] as was done in [23]. Then, dimensionality reduction techniques given in the Appendix could be used to reduce the total number of modes. Subject specific head models can be used to design fdTMS coils. However, accurately representing the stimulated region within the geometrically complex gyrus requires an excessive amount of sample points, and the optimization using our MILP approach becomes computationally intractable. A future framework for subject-specific coil design accounting for gyrification could leverage active subspace methods [41] for determining a minimal set of modes that spans the fdTMS coil design space. This will result in significant dimensionality reduction because of the diffusive relation between coil current distribution and cortical E-field. We anticipate that for practical energy levels this dimensionality reduction will enable the use of global optimization techniques like random search and genetic algorithms to find the optimum coil designs.

Note that in computationally optimizing TMS coils the E-field has to be sampled with finite spatial resolution. The spatial resolution of the meshes used in this study was on the order of 1 mm; as such we captured variations in the E-field in that range. Since typical ranges for spread are on the order of cm^2^, we do not expect further significant improvements in depth–focality beyond what was reported here. Moreover, the finite sampling rate quantizes the possible improvements in depth–focality trade-off that the optimization can observe. In turn, this causes intervals of energy levels for which the optimization achieves the same spread for a given depth; this results in slight energy suboptimality for coils at high energy levels. Nevertheless, these small energy suboptimalities can be suppressed by optimizing on a denser mesh or alternatively running an energy minimization on the coil like the one proposed in [25].

Consistent with previous design studies [12, 23], the energy required by commercial TMS coils can be reduced significantly while preserving the field shape. Furthermore, the fdTMS designs achieved significantly increased focality and depth even at energy levels of existent coil drivers. Energy efficiency of fdTMS coils could be improved further by introducing a ferromagnetic core directly above them [42-44], thereby enabling better focality vs. energy trade-offs. However, ferromagnetic cores introduce a nonlinearity in the relationship between the coil current and the resultant magnetic field that can distort the induced E-field. The degree to which this affects the coil focality and depth characteristics is expected to be limited since the core shapes the magnetic field mostly on the backside of the coil, away from the subject’s head [20]. In any case, the effects of a ferromagnetic core can be evaluated with simulations [18, 20] and potentially linked to the optimization framework in an iterative process.

Once the fdTMS coil windings are determined, the coil can be manufactured with a high level of automation. For example, the coil can be made by 3D-printing a plastic former with grooves in which copper wire is inserted [12, 23, 25]. Alternatively, planar coils can be implemented with a stack of printed circuit boards with heavy copper traces following the winding design [45].

## CONCLUSIONS

We estimated the limit of TMS coil focality as a function of stimulation depth. It was shown that existing coil designs do not reach this limit. For a given maximum depth of stimulation, spread can be theoretically reduced about two-fold compared to conventional coils. A substantial fraction of these improvements can be achieved with feasible energy requirements. These results appear to be the first systematic advancement in the depth–focality trade-off of TMS coils since the introduction of the figure-8 coil three decades ago [4], and likely represent the fundamental physical limit.

## ACKNOWLEDGEMENT

Results from this work were presented in abstract form at the 4^th^ Annual BRAIN Initiative Investigators Meeting, Rockville, MD, USA, April 9–11, 2018. We thank Dr. Risto Ilmoniemi and Dr. Lari Koponen for reading and providing helpful suggestions on this manuscript.

## APPENDIX Adaptive Refinement

Here we describe an iterative and adaptive scheme for solving optimization problem defined in Eq. (21) of the paper. It is assumed that we have a tetrahedron mesh of the brain and E-field values are available everywhere inside it. The numerical experiments indicate that to obtain an optimal design, it is only necessary to sample the E-field on the surface of the tetrahedral mesh; thus, we only consider field samples on it. Furthermore, the sampling scheme described here is meant to be a low-fidelity scheme that sufficiently approximates *V*(*λ*) in a way that coils can be ranked to enable optimization while maintaining computational tractability. After optimization, *V* (*λ*) is computed to high accuracy by sampling the E-field on all tetrahedron centers and using the tetrahedron volume as sample weights.

**Figure A1.**
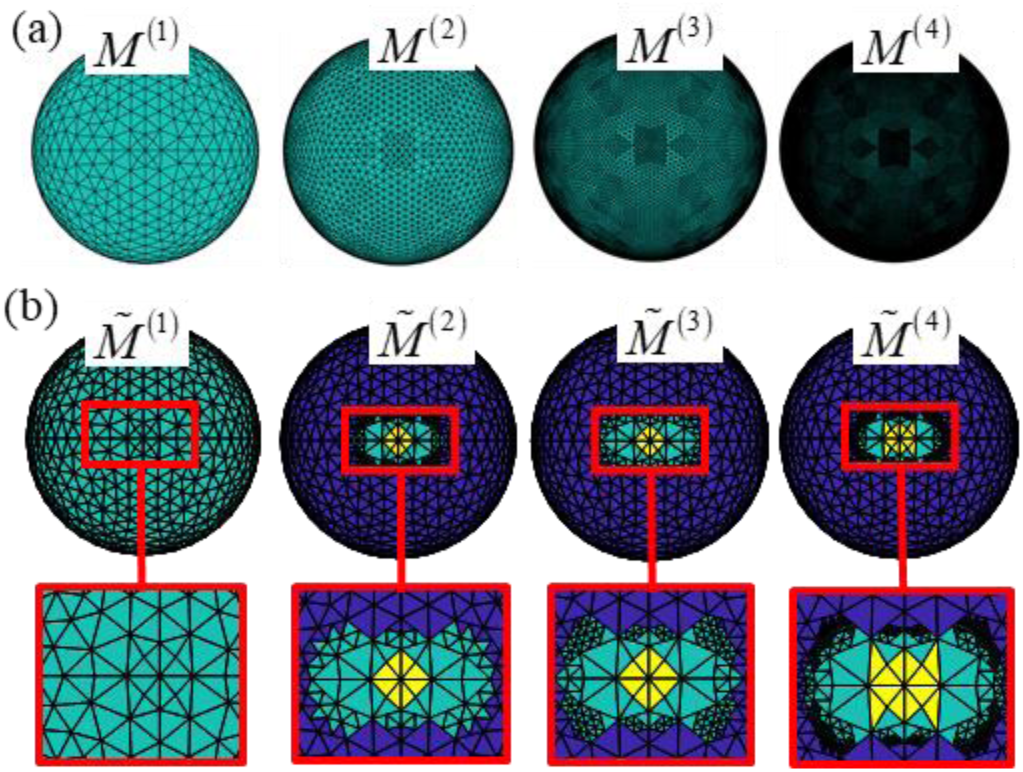
Mesh family sets. (a) Family set of meshes for the spherical head model with increasing density. (b) Family set of meshes for a sample optimization. Macro-cells that are included as optimization variables (cyan), assumed stimulated (yellow), and not-stimulated (blue).

Starting from a coarse representation of the cost function in Eq. (22), the adaptive procedures solves a series of MILP problems each time refining regions where the optimal design generates E-fields near threshold. This results in a discretization that properly captures the transition from stimulated region to non-stimulated region with minimal number of quadrature points. To do the above, the adaptive sampling procedure leverages a hierarchical family set of *L* brain boundary meshes. The meshes are numbered *i* = 1, 2, …, *L* and each mesh *M*^(*i*)^ is composed of *N*^(*i*)^ macro-cells generated by combining a number adjacent triangles of the brain boundary mesh. Macro-cells of mesh *M*^(*L*)^ are the triangles of the brain boundary mesh (i.e. *M*^(*L*)^ is the original mesh). Starting from mesh *M*^(*L*)^, level-(*i* − 1) mesh *M*^(*i*−1)^ will be formed by combining the adjacent macro-cells of the level-*i* mesh *M*^(*i*)^ so that their average area is about four times the average area of groups of level-*i*. Correspondingly, each macro-cell *M*^(*i*)^ is a combination of macro-cells of mesh *M*^(*i*+1)^; these macro-cells are called its children. Figure A1(a) depicts mesh family set of the sphere head model. The sphere head family mesh set was generated by barycentric refinement of a sphere mesh and the SimNIBS head mesh family set was generated via an oct-tree procedure [46]. For each mesh *M*^(*i*)^, each macro-cell is assigned an E-field
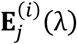
(where *j* = 1, …,*N*^(*i*)^) equal to the average total E-field on it when exposed to the primary E-field of **I**(**r**; *λ*). Correspondingly, the family set of meshes forms a hierarchy from coarsest (*M*^(1)^) to finest (*M*^(*L*)^) of E-field representations.

Note that each optimization macro-cell that satisfies
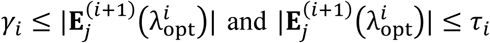
is assumed to be stimulated and not-stimulated, respectively, by
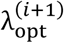
and their slack variables are not included in the optimization. A sample hierarchy of meshes *M͠*^(1)^ to *M͠*^(4)^ are shown in figure A1(b). Finer regions of the mesh *M͠*^(4)^ correspond to regions that are exposed to an E-field near *E*_TH_; this enables accurate stimulated region representation with few samples. In our optimizations, *L* = 4 and values for parameters *χ_i_*, *β_i_*, *γ_i_* and *τ_i_* are given in table A1.

**Table A1.**
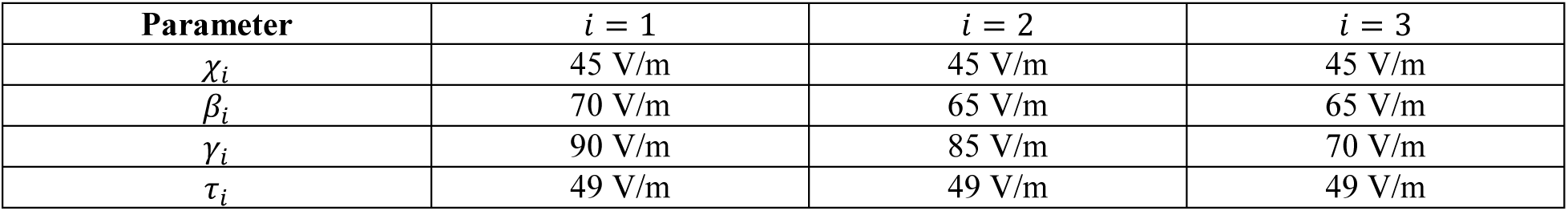
Values of parameters *χ_i_*, *β_i_*, *γ_i_* and *τ_i_* of adaptive sampling procedure at each level *i*.

| Parameter | $i = 1$ | $i = 2$ | $i = 3$ |
| --- | --- | --- | --- |
| $\chi_i$ | 45 V/m | 45 V/m | 45 V/m |
| $\beta_i$ | 70 V/m | 65 V/m | 65 V/m |
| $\gamma_i$ | 90 V/m | 85 V/m | 70 V/m |
| $\tau_i$ | 49 V/m | 49 V/m | 49 V/m |

## Mode Preprocessing

Here the approach used for converting a set of linearly independent modes **M**_*i*_(**r**) (where *i* = 1,2,…, *N*) into energy orthonormal reduced basis of modes **M̃̃**_*i*_(**r**) (where *i* = 1,2,…,*Ñ̃*) is outlined. First, modes **M**_*i*_(**r**) are replaced by energy orthonormal modes **M̃**_*i*_(**r**) that span the same space of surface currents. Second, the basis modes **M̃**_*i*_ (**r**) are replaced with a reduced basis set of modes **M̃̃**_*i*_ (**r**) that only span currents that efficiently couple into the head.

Current modes **M̃**_*i*_(**r**) are determined from the singular value decomposition (SVD) of the mutual inductance matrix **L** = **USV^T^**. Because **L** is positive definite, **U** = **V** and **L** = **VDV^T^**. Energy orthonormal modes are

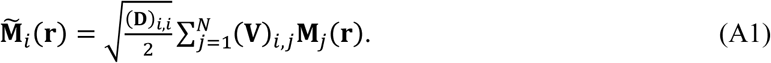

Many of the current distributions spanned by **M̃**_*i*_(**r**) are energy inefficient (i.e. require orders of magnitude more energy than what is delivered to the brain and the electromagnetic coupling between these modes and the brain is low). A second transformation is used to modify the basis consisting of modes **M̃**_*i*_(**r**) (where *i* = 1,2,…, *N*) to a reduced basis consisting of modes **M̃̃**_*i*_ (**r**) (where *i* = 1,2,…, *Ñ̃*) that do not span energy inefficient currents. The transformation is outlined in the next few paragraphs and relies on auxiliary energy optimal current basis modes for generating E-field in the brain. First, we define an auxiliary energy optimal current basis mode set and a matrix that links coefficients for a design in the span of basis set **M̃**_*i*_ (**r**) to coefficients of energy optimal equivalent ones. Then, this matrix is used to define new reduced basis set **M̃̃**_*i*_(**r**). The procedure can be done numerically for arbitrary head and coil positioning, but for notational brevity we assume that the coil is placed 5 mm above the apex of the head, the spherical head model is used, and the spherical model is centered about the coordinate origin.

Consider energy optimal current mode **H**_*i*_(**r**) that generates the same E-field in a spherical brain region of *r_b_* radius as their corresponding mode **M̃**_*i*_(**r**). In [12] it was shown that **H**_*i*_(**r**) will reside on the surface of the *r_b_* sphere and

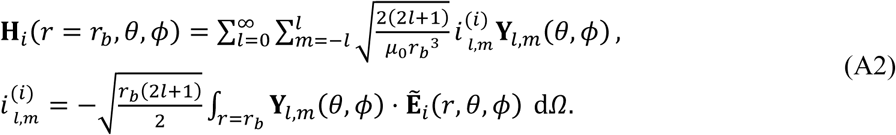

Here **Ẽ**_*i*_(**r**) is the E-field generated by **M̃**_*i*_(**r**), **Y**_*l*,*m*_(*θ*,*ϕ*) is a vector spherical harmonic as defined in [12], the integration is performed over the entire spherical surface, and *dΩ* = *r*^2^sin(*θ*)*dθdϕ*. Furthermore, each mode **H**_*i*_(**r**) has energy

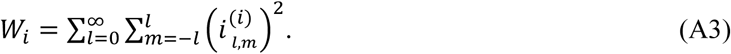

A current distribution

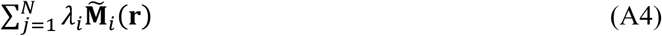

resulting from a vector of weights *λ* = (*λ*_1_, *λ*_2_,…, *λ_N_*) will have corresponding minimum energy current

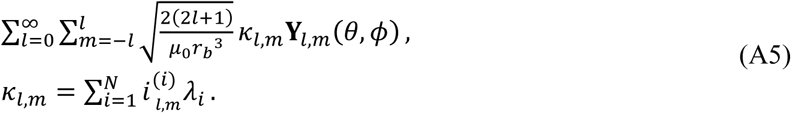

Assuming that we only compute the coefficients up to terms *l* = *l*_max_ (we choose *l*_max_ = 20) the coefficients are related by a matrix vector multiplication

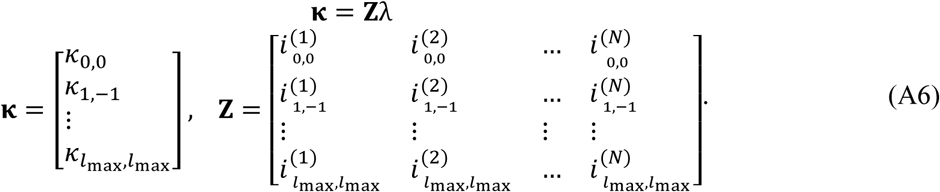

The matrix **Z** relates currents spanned by modes **M̃**_*i*_(**r**) to their minimum energy coil spanned by modes **H**_*i*_(**r**). Furthermore, Eq. (A6) implies that the energy of the lowest energy surface current generating the same field as *λ* will be ∥**κ**∥^2^ = ∥**Z***λ*∥^2^. Correspondingly, right singular column vectors **ṽ**_1_ (*i* = 1,2, …,*N*) of **Z** result in energy orthonormal modes with surface currents

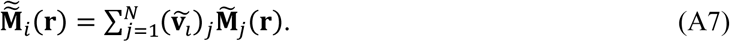

Furthermore, for each mode **M̃̃**_*i*_(**r**), the singular value *S_i_* corresponding to **ṽ**_*l*_ is equal to the square root of its energy efficiency. Only modes **M̃̃**_*i*_(**r**) that have an energy efficiency above *ε_eff_* = 10^−4^
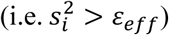
are included in the optimization.

## REFERENCES

[1] O’Reardon J P, Solvason H B, Janicak P G, Sampson S, Isenberg K E, Nahas Z, McDonald W M, Avery D, Fitzgerald P B and Loo C 2007 Efficacy and safety of transcranial magnetic stimulation in the acute treatment of major depression: a multisite randomized controlled trial Biological psychiatry 62 1208–16

[2] Lipton R B, Dodick D W, Silberstein S D, Saper J R, Aurora S K, Pearlman S H, Fischell R E, Ruppel P L and Goadsby P J 2010 Single-pulse transcranial magnetic stimulation for acute treatment of migraine with aura: a randomised, double-blind, parallel-group, sham-controlled trial The Lancet Neurology 9 373–80

[3] Picht T, Krieg S M, Sollmann N, Rösler J, Niraula B, Neuvonen T, Savolainen P, Lioumis P, Mäkelä J P and Deletis V 2013 A comparison of language mapping by preoperative navigated transcranial magnetic stimulation and direct cortical stimulation during awake surgery Neurosurgery 72 808–19

[4] Ueno S, Tashiro T and Harada K 1988 Localized stimulation of neural tissues in the brain by means of a paired configuration of time ^−^ varying magnetic fields J. Appl. Phys. 64 5862–4

[5] Chunye R, Tarjan P P and Popovic D B 1995 A novel electric design for electromagnetic stimulation-the Slinky coil IEEE Trans. Biomed. Eng. 42 918–25

[6] Kai-Hsiung H and Durand D M 2001 A 3-D differential coil design for localized magnetic stimulation IEEE Trans. Biomed. Eng. 48 1162–8

[7] Davey K R and Riehl M 2006 Suppressing the surface field during transcranial magnetic stimulation IEEE Trans. Biomed. Eng. 53 190–4

[8] Dong-Hun K, Georghiou G E and Won C 2006 Improved field localization in transcranial magnetic stimulation of the brain with the utilization of a conductive shield plate in the stimulator IEEE Trans. Biomed. Eng. 53 720–5

[9] Lu M 2009 Calculating the electric field in real human head by transcranial magnetic stimulation with shield plate J. Appl. Phys. 105 07B322

[10] Hernandez-Garcia L, Hall T, Gomez L and Michielssen E 2010 A numerically optimized active shield for improved transcranial magnetic stimulation targeting Brain stimulation 3 218–25

[11] Gomez L, Hernandez L, Grbic A and Michielssen E 2010 A simulation of focal brain stimulation using metamaterial lenses. In: Antennas and Propagation Society International Symposium (APSURSI), 2010 IEEE, pp 1–4

[12] Koponen L M, Nieminen J O and Ilmoniemi R J 2015 Minimum-energy coils for transcranial magnetic stimulation: application to focal stimulation Brain stimulation 8 124–34

[13] Ruohonen J and Ilmoniemi R 1998 Focusing and targeting of magnetic brain stimulation using multiple coils Medical and Biological Engineering and Computing 36 297–301

[14] Ho S L, Guizhi X, Fu W N, Qingxin Y, Huijuan H and Weili Y 2009 Optimization of Array Magnetic Coil Design for Functional Magnetic Stimulation Based on Improved Genetic Algorithm IEEE Trans. Magn. 45 4849–52

[15] Han B H, Chun I K, Lee S C and Lee S Y 2004 Multichannel magnetic stimulation system design considering mutual couplings among the stimulation coils IEEE Trans. Biomed. Eng. 51 812–7

[16] Gomez L J, Hernandez-Garcia L, Grbic A and Michielssen E 2013 Single-source multi-coil transcranial magnetic stimulators for deep and focused stimulation of the human brain. In: Radio Science Meeting (Joint with AP-S Symposium), 2013 USNC-URSI: IEEE) pp 8-

[17] Roth Y, Zangen A and Hallett M 2002 A Coil Design for Transcranial Magnetic Stimulation of Deep Brain Regions Journal of Clinical Neurophysiology 19 361–70

[18] Deng Z-D, Lisanby S H and Peterchev A V 2014 Coil design considerations for deep transcranial magnetic stimulation Clinical Neurophysiology 125 1202–12

[19] Rastogi P, Lee E G, Hadimani R L and Jiles D C 2017 Transcranial Magnetic Stimulation-coil design with improved focality AIP Advances 7 056705

[20] Deng Z-D, Lisanby S H and Peterchev A V 2013 Electric field depth–focality tradeoff in transcranial magnetic stimulation: simulation comparison of 50 coil designs Brain stimulation 6 1–13

[21] Gomez L, Cajko F, Hernandez-Garcia L, Grbic A and Michielssen E 2013 Numerical analysis and design of single-source multi-coil TMS for deep and focused brain stimulation IEEE Trans. Biomed. Eng. 60 2771–82

[22] Goetz S M, Weyh T, Afinowi I A A and Herzog H G 2014 Coil Design for Neuromuscular Magnetic Stimulation Based on a Detailed 3-D Thigh Model IEEE Trans. Magn. 50 1–10

[23] Koponen L M, Nieminen J O, Mutanen T P, Stenroos M and Ilmoniemi R J 2017 Coil optimisation for transcranial magnetic stimulation in realistic head geometry Brain Stimulation

[24] Sánchez C C, Rodriguez J M G, Olozábal Á Q and Blanco-Navarro D 2016 Novel TMS coils designed using an inverse boundary element method Physics in Medicine and Biology 62 73

[25] Wang B, Shen M R, Deng Z-D, Smith J E, Tharayil J J, Gurrey C J, Gomez L J and Peterchev A V 2018 Redesigning existing transcranial magnetic stimulation coils to reduce energy: application to low field magnetic stimulation Journal of neural engineering

[26] Peeren G N 2003 Stream function approach for determining optimal surface currents Journal of Computational Physics 191 305–21

[27] Huang Y, Parra L C and Haufe S 2016 The New York Head—A precise standardized volume conductor model for EEG source localization and tES targeting NeuroImage 140 150–62

[28] Jackson J D 1975 Electrodynamics: Wiley Online Library)

[29] Peterchev A V, Deng Z-D and Goetz S M 2015 Brain Stimulation: John Wiley & Sons) pp 165–89

[30] Guadagnin V, Parazzini M, Fiocchi S, Liorni I and Ravazzani P 2016 Deep transcranial magnetic stimulation: modeling of different coil configurations IEEE Trans. Biomed. Eng. 63 1543–50

[31] Opitz A, Windhoff M, Heidemann R M, Turner R and Thielscher A 2011 How the brain tissue shapes the electric field induced by transcranial magnetic stimulation Neuroimage 58 849–59

[32] Jin J-M 2014 The finite element method in electromagnetics: John Wiley & Sons)

[33] Ylä-Oijala P and Taskinen M 2003 Calculation of CFIE impedance matrix elements with RWG and n× RWG functions IEEE Trans. Antennas Propagat. 51 1837–46

[34] Freund R W 1993 A transpose-free quasi-minimal residual algorithm for non-hermitian linear systems SIAM J. Sci. Stat. Comput. 14 470–82

[35] Udell M and Boyd S 2013 Maximizing a sum of sigmoids Optimization and Engineering

[36] MATLAB Optimization Toolbox Release 2017b. (Natick, MA: The MathWorks, Inc.)

[37] Teanby N A 2006 An icosahedron-based method for even binning of globally distributed remote sensing data Computers & Geosciences 32 1442–50

[38] Seiler M C and Seiler F A 1989 Numerical recipes in C: the art of scientific computing Risk Analysis 9 415–6

[39] Thielscher A, Antunes A and Saturnino G B 2015 Field modeling for transcranial magnetic stimulation: A useful tool to understand the physiological effects of TMS? In: 2015 37th Annual International Conference of the IEEE Engineering in Medicine and Biology Society (EMBC), pp 222–5

[40] Vecchi G 1999 Loop-star decomposition of basis functions in the discretization of the EFIE IEEE Trans. Antennas Propagat. 47 339–46

[41] Constantine P G, Dow E and Wang Q 2014 Active subspace methods in theory and practice: applications to kriging surfaces SIAM J. Sci. Comput. 36 A1500–A24

[42] Davey K and Epstein C M 2000 Magnetic stimulation coil and circuit design IEEE Trans. Biomed. Eng. 47 1493–9

[43] Lorenzen H W and Weyh T 1992 Practical application of the summation method for 3-D static magnetic field calculation of a setup of conductive and ferromagnetic material IEEE Trans. Magn. 28 1481–4

[44] Epstein C M and Davey K R 2002 Iron-core coils for transcranial magnetic stimulation Journal of Clinical Neurophysiology 19 376–81

[45] Goetz S M, Smith E J, Gomez L J and Peterchev A V 2018 Coil design for quiet transcranial magnetic stimulation (qTMS). In: 2018 BRAIN Initiative Principal Investigators Meeting, (Rockville, MD)

[46] Chew W C, Jin J-M, Lu C-C, Michielssen E and Song J M 1997 Fast solution methods in electromagnetics IEEE Trans. Antennas Propagat. 45 533–43

